# Predicting evolution using frequency-dependent selection in bacterial populations

**DOI:** 10.1101/420315

**Authors:** Taj Azarian, Pamela P Martinez, Brian J Arnold, Lindsay R Grant, Jukka Corander, Christophe Fraser, Nicholas J Croucher, Laura L Hammitt, Raymond Reid, Mathuram Santosham, Robert C Weatherholtz, Stephen D Bentley, Katherine L O’Brien, Marc Lipsitch, William P Hanage

## Abstract

Predicting how pathogen populations will change over time is challenging. Such has been the case with *Streptococcus pneumoniae*, an important human pathogen, and the pneumococcal conjugate vaccines (PCVs), which target only a fraction of the strains in the population. Here, we use the frequencies of accessory genes to predict changes in the pneumococcal population after vaccination, hypothesizing that these frequencies reflect negative frequency-dependent selection (NFDS) on the gene products. We find that the standardized predicted fitness of a strain estimated by an NFDS-based model at the time the vaccine is introduced enables to predict whether the strain increases or decreases in prevalence following vaccination. Further, we are able to forecast the equilibrium post-vaccine population composition and assess the invasion capacity of emerging lineages. Overall, we provide a method for predicting the impact of an intervention on pneumococcal populations with potential application to other bacterial pathogens in which NFDS is a driving force.

## Introduction

Human interventions perturb microbial populations in many ways. Most obviously, the use of antibiotics or vaccines that target some strains and not others provide opportunities for new strains to emerge and become established. Examples include vaccines for antigenically diverse human pathogens like influenza, *Neisseria meningitidis, Haemophilus influenzae, Streptococcus pneumoniae*, and human papillomavirus [1–3]. Predicting these changes is a central goal of population genomic and evolutionary studies of pathogens [4–7]. For bacteria in particular, detailed predictions of how a population will respond to a selective pressure are challenging. Models that specify how mutations with a given fitness change in frequency over time are often hard to apply in practice, as we typically do not know in advance important parameters such as the fitness value of particular alleles or how this is affected by their frequency (frequency-dependent selection) or genetic background (epistasis) [8,9].

Ongoing efforts to control disease caused by *Streptococcus pneumoniae* (the pneumococcus), a colonizer of the human nasopharynx and a cause of pneumonia, bacteremia, meningitis, and otitis media, underscore the difficulties of predicting changes after introduction of a vaccine [10]. Pneumococcal conjugate vaccines (PCVs) target only a fraction of this antigenically diverse species, which contains over 90 distinct serotypes [11]. Following widespread introduction of PCVs, non-vaccine serotypes (NVT) benefitted from the removal of their vaccine-serotype (VT) competitors and became more common in carriage and disease, with the gains from reducing VT disease partly offset by increases in NVT disease [12–14]. These changes in the pathogen population varied by location and were not fully appreciated until retrospective analysis [15–17].

Our recent study of pneumococcal carriage isolates collected before and after PCV7 vaccine introduction in the southwest US [17] illustrates the complexity in post-vaccine population dynamics, echoing findings from other studies. Pneumococcal populations contain multiple ‘sequence clusters’ which are closely related lineages, defined on the basis of sequence variation in loci present among all isolates (i.e., the core genome) [18]. We henceforth use the term *strains* to refer to these lineages/sequence clusters. Variation in genome content due to horizontal gene transfer is a hallmark of prokaryotes; therefore, in addition to the core genome, we can define the accessory genome, as those genes not found in all isolates in the sample [19,20]. Consistent with their close phylogenetic relatedness in terms of core genome sequence variation, each strain we identify is comprised of isolates that are fairly homogeneous – but not completely so – in the presence/absence of accessory genes as well as phenotypic properties such as serotype and antibiotic resistance [21].

Previous work showed that post-vaccine success of pneumococcal strains may depend on the accessory genome [22,23]. In many bacteria, this can be a large fraction of the total number of genes found in a species (i.e., the ‘pangenome’) [24,25]. A population genomic study of pneumococci in Massachusetts children found that vaccination had remarkably little effect, after six years, on the overall frequencies of individual accessory genes (defined as clusters of orthologous genes or COGs) [23]. Despite the fact that nearly half the pre-vaccine population had serotypes targeted by the vaccine, only two of >3000 loci in the accessory genome significantly decreased in frequency 6 years post-introduction, and none increased [23]. More recently, a geographically diverse sample of pneumococcal genomes showed that while the distribution of strains varied widely across the globe, the proportion of isolates in each sample containing each individual accessory gene was highly consistent across locations [22]. Where vaccine was introduced, accessory gene frequencies were perturbed by the removal of vaccine types but trended back toward their pre-vaccine frequencies over time [17,22,26]. Negative frequency-dependent selection (NFDS) was proposed as the mechanism by which the frequencies of loci were restored after vaccine introduction [22]. NFDS is a type of balancing selection, which maintains diversity by favoring variants when rare, but exacting a cost when they become common, such that the frequency of the variant stabilizes at intermediate values, or in some instances result in frequency oscillations [9]. Examples of mechanisms produced by NFDS include host immunity and bacteriophage predation, and as such, balancing selection is recognized as a key contributor to population composition and diversity [27,28]. Among pneumococci, similar processes have been proposed to explain the co-existence of multiple serotypes [29] and vaccine-induced metabolic shifts [30].

Here, we present flexible, easily computable statistics that estimate the fitness of any strain using the contents of its accessory genome as a proxy for how it will be affected by NFDS, dependent on the frequencies of other strains in the population, and specifically of the accessory genes they carry. Even though we do not know the specific loci under selection or the mechanism involved, we are able to make predictions about the composition of a population as well as predict the fitness of any strain in any population, whether or not it has yet appeared in that population. Overall, this predictive model offers a way to study population processes and the response to interventions.

## Results

In the sample of 937 pneumococcal isolates comprised of 35 strains from the southwest US, we observed a sharp decline in PCV7-VT strains following vaccination (Figure 1 and S1 Figure). VT strains were subsequently replaced by NVT strains, including two emergent NVT strains that had not been observed pre-vaccination, although they were present during the same time period in a related carriage dataset from Massachusetts [17,23]. We first show that there was considerable deviation from the null expectation that NVT strains would increase in prevalence *pro rata* to their pre-vaccine frequency; the most common NVT strains before vaccination were not necessarily the most prevalent 12 years afterwards (Figure 1A). In particular we find 13 of 35 strains deviated significantly from the prevalence expected under a null *pro rata* model; 9 were more common than expected and 4 less common, annotated with plus and minus signs, respectively, in Figure 1B. The impact of vaccination on individual NVT strains was hence not easily predictable. Consequently, public health authorities and vaccine manufacturers have had to rely on post-vaccine surveillance to estimate the next epidemiologically important lineage and determine subsequent vaccine formulations. At best, this uncertainty reduces the population impact of vaccination; at worst, it could unintentionally increase the prevalence of virulent or antibiotic resistant lineages [31].

**Figure 1.**
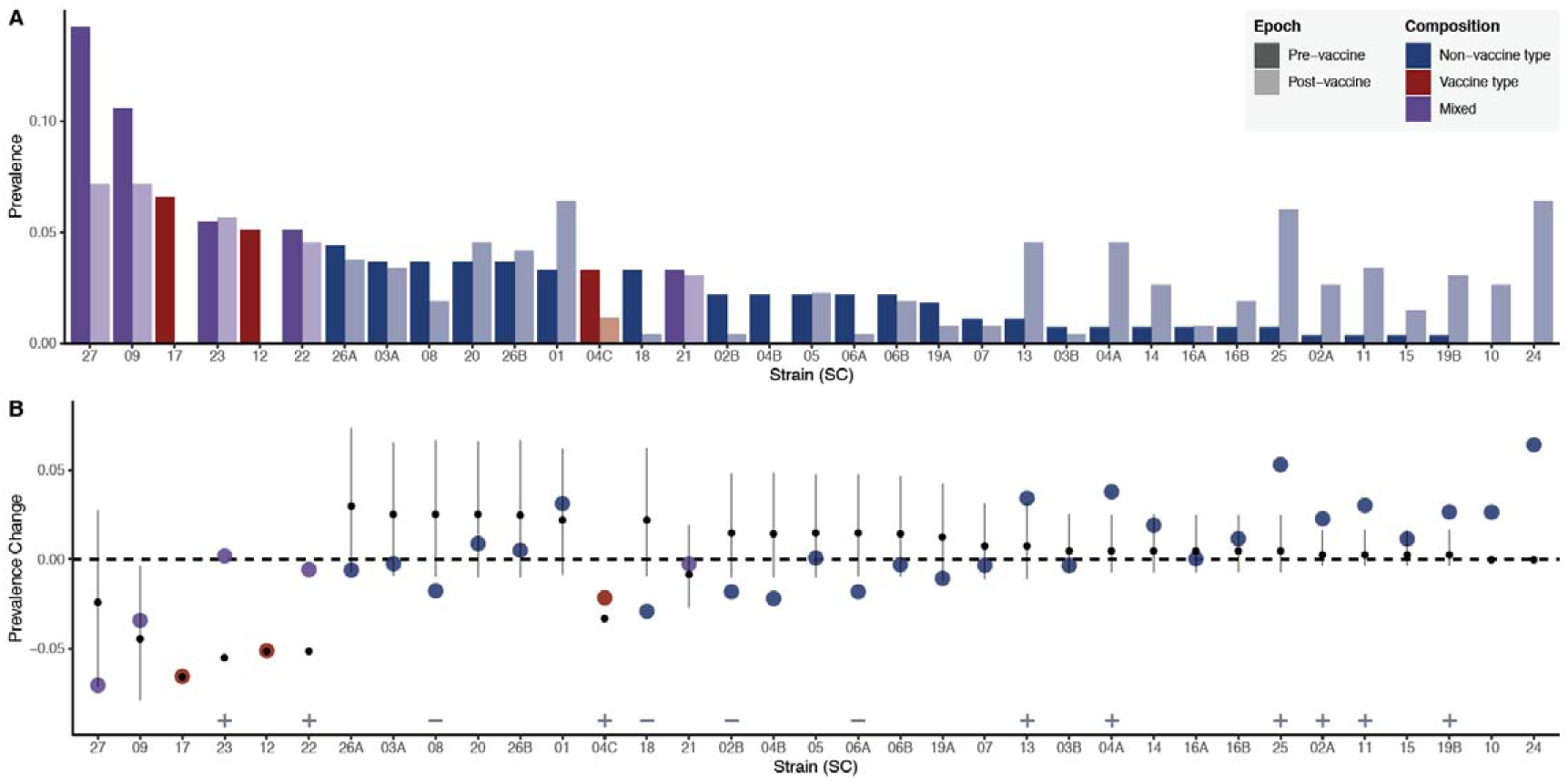
A.) Pre-vaccine to post-vaccine change in prevalence of strains (SCs). Strains are ordered from highest to lowest pre-vaccine prevalence. **B.) Observed prevalence change calculated as post-vaccine frequencies minus pre-vaccine frequencies**. Changes in prevalence are compared to that expected under a *pro rata* null model (i.e., not using the predictive methods in this paper). Observed changes in prevalence are represented by points colored by the serotype composition of the strain: non-vaccine serotype (NVT) only, PCV7 vaccine-serotype (VT) only, and mixed VT and NVT (VT-NVT). The point and whiskers show the prevalence change expected if all VT strains were removed and NVT increased proportional to their pre-vaccine prevalence – i.e., in a null model of *pro rata* increase where only the VT strains were removed and all NVT strains increased equally in proportion to their pre-vaccine prevalence. The dot is the median, and the whiskers give the 2.5% and 97.5% quantiles of predicted changes under the null model using 10,000 bootstraps from pre-vaccine samples. Significant differences between the changes in prevalence from the *pro rata* model and the observed data are denoted with plus and minus signs specifying strains that were significantly more (n=9) or less (n=4) common, respectively. Among the most successful were strains that contained both VT and NVT isolates (SC-22 and SC-23) whose NVT component included serotypes 6C, 15C, and 35B, as well as SC-24 and SC-25, which were dominated by the NVT serotypes 23A and 15C, respectively. SC-27 is polyphyletic, comprised of an aggregate of strains that are at low frequency in the overall population. Compared to strains comprised of solely NVT isolates, those with mixed NVT-VT had marginally higher risk differences, indicating greater success than expected under the null model (*β*=0.03, SE=0.015, F(1,29)=3.67, p=0.06). Two strains that emerged during the study period (SC-10 and SC-24) were not included in this analysis as they were not present at the first time point.

Having documented that there were strains that increased significantly more or less than their pre-vaccine frequency would indicate, we sought to define a parsimonious predictive algorithm based on NFDS that could account for these changes. We hypothesized that evolutionary dynamics could be predicted on the premise that after perturbation by vaccine, strains characterized by accessory genomes that could best restore the pre-perturbation accessory-gene-frequency equilibrium would have the highest fitness and therefore increase in prevalence disproportionately. To this end, we implemented a deterministic model using the replicator equation to calculate the fitness of a strain based on its accessory genome, using vaccination as an example of perturbation [32–34] (equation 1).

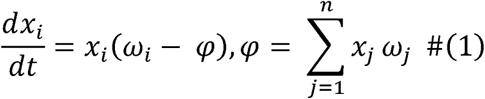

Under this formulation, *x*_*i*_ denotes the frequency of strain *i* (*i* = {1,…, *n*}), *n* is the total number of strains, *ω*_*i*_ denotes the fitness of strain *i* (adapted from Ref. [22]), and *φ* is the average population fitness. The difference (*ω* − *φ*) is a standardized predicted fitness, and the fitness vector *ω* is defined as the product of matrix **K** whose element *k*_*i,l*_ is a value between 0 and 1 for the frequency of accessory gene *l* in strain *i*, and the vector (*e* − *f*) whose *I*^*th*^ element is the difference between the pre-vaccine frequency *e*_*l*_ and *f*_*l*_, which is the gene’s expected frequency post-vaccination, based on removing the VTs from the pre-vaccine population, of each accessory gene *I* (equation 2). Intuitively, the vector (*e* − *f*) represents the vacancy that vaccination produces in the population in terms of the accessory loci it removes, and *ω*_*i*_ quantifies the ability of strain *i* to fill that gap. In contrast with previous work [22], we do not define carrying capacity or migration rates, requiring only knowledge of the accessory gene frequencies at equilibrium and which strains they are associated with; these quantities can be estimated from a population survey prior to the perturbation of interest. We assume that the impact of recombination on the accessory genome is negligible over the relatively short time period we study here.

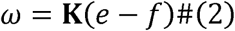

Using simulated data, we first assessed the ability of a strain’s standardized predicted fitness (*ω* − *φ*) (for brevity we drop the modifier “standardized” hereafter) to predict the direction of its change in frequency, based on its ability to resolve the vaccine-induced perturbation (Figure 2). Note that this predicted fitness uses only data available before vaccine rollout. Using this model, we show that in simulations, the predicted fitness is consistent with the direction of a simulated strain’s adjusted prevalence change (i.e. changes in prevalence minus what would be expected if all NVT strains increased by the same proportion from their pre-vaccine prevalence) 92.8% of cases, independent of the initial pre-vaccine frequency (Figure 2B). Next, we asked whether this approach could predict the post-vaccine composition of an actual pneumococcal population, and specifically the relative contribution of each strain to serotype replacement. For each strain present before vaccine introduction, we used the accessory genome to calculate the fitness following the removal of vaccine types. We identified 2,371 genes that were present in between 5% and 95% of isolates. In this data set, we found the predicted fitness value was significantly and positively correlated with the observed prevalence change (Adjusted R^2^=0.41, p<<0.001, Figure 3A). Further, the trajectory following vaccination, whether increasing or decreasing in frequency, was accurately predicted for 28 of the 31 tested strains identified in the sample, as indicated by the upper right and lower left quadrants of Figure 3A. Strains with a positive prevalence change had substantially higher predicted fitness than those with a negative one (mean fitness of strains that increased vs. decreased 6.4 vs. −2.4; 95% CI of the difference: 5.0-12.5, p<0.001).

**Figure 2.**
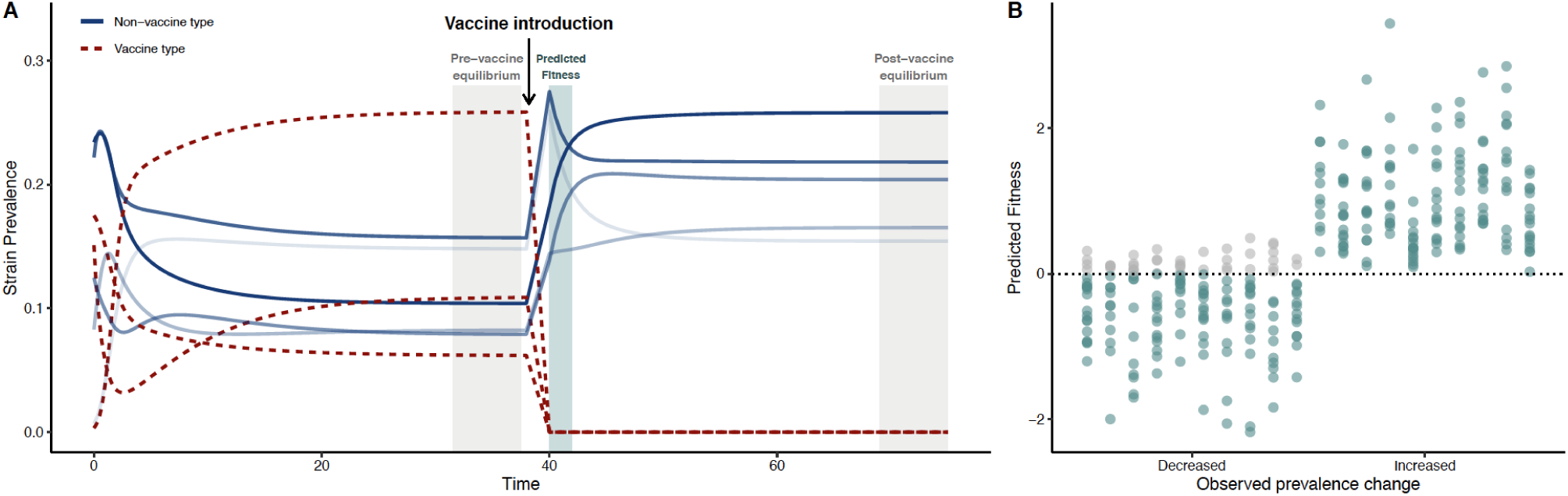
A.) Conceptual diagram for simulations. Descriptive representation of the strain prevalence at different stages relative to vaccine introduction: pre-vaccine equilibrium, vaccine introduction, and post-vaccine equilibrium. We modeled a population of VT and NVT strains (represented as unique genotypes with alleles 1 or 0 at a locus, denoting the presence or absence of a single accessory locus) and simulated the removal of VT genotypes, following the post-vaccine population to equilibrium (details in methods). In this illustrative figure, eight strains are shown, with their prevalence in the population evolving over time. The system is allowed to evolve until it reaches a steady state (‘pre-vaccine equilibrium’). Three strains were then targeted to mimic a vaccine introduction, which removes them from the system. The predicted fitness was then estimated from the period just after the vaccine introduction, when the population has been depleted of VT but relative prevalence of NVT has not changed – a quantity that can be calculated from pre-vaccine data alone. Finally, the system reaches a second steady state (‘post-vaccine equilibrium’). Different shades of blue represent the rank of the strain frequencies in the post-vaccine equilibrium. **B.) Simulation results**. Comparison of the direction of prevalence change of strains from pre-to post-vaccine using simulated data and predicted fitness from these simulated data. For these 10 replicate simulations, 2,371 accessory loci and 35 randomly chosen strains were simulated, including three VT genotypes. For each replicate, the pre-vaccine equilibrium frequencies of the 2,371 accessory loci were varied. Final prevalence of strains were obtained by quadratic programing, and prevalence change for each NVT strain was calculated as post-vaccine prevalence minus pre-vaccine prevalence, in both cases with all NVT strains summing to 100%. Each column in the decreased and increased category represents the results from one simulation (i.e., the first column in the decreased category corresponds to the first column in the increased category and the dots sum to 32). The predicted fitness of the strain accurately predicts the direction of the prevalence change in 92.8% of cases (teal dots). Grey dots represent instances where the direction of the prevalence change was not predicted correctly in the simulation.

**Figure 3.**
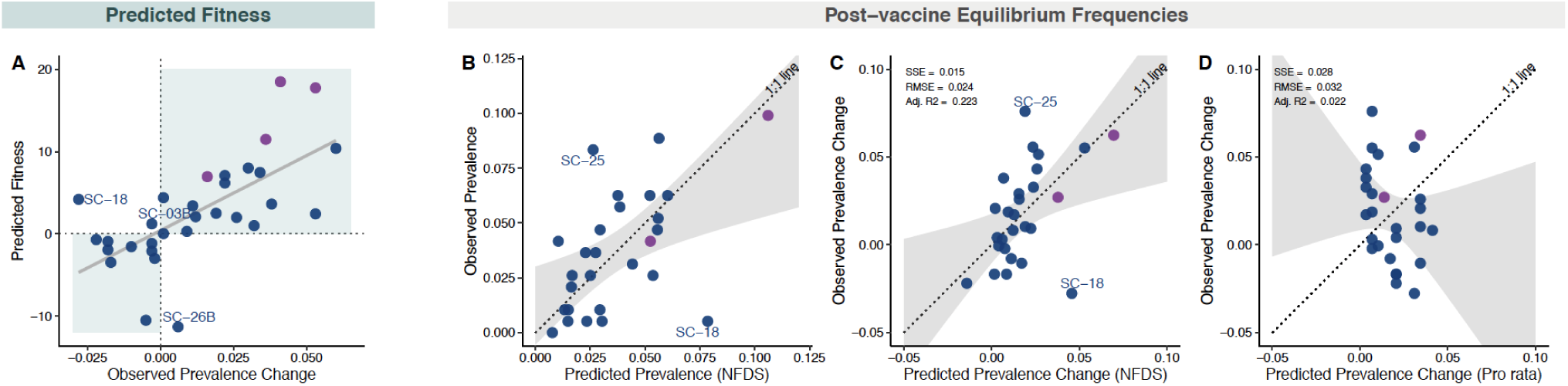
A.) Relationship between predicted fitness and observed prevalence change from pre- to post-vaccine among 31 strains, in each case summing to 100%. Prevalence change was calculated as post-vaccine frequencies minus pre-vaccine frequencies. Predicted fitness was calculated using data solely from the pre-vaccine sample, with the exceptions of strains for which there were no non-vaccine serotype (NVT) isolates present in the sample before the introduction of PCV7 (n=4). For those strains, data were imputed from the time point during which they were first observed. Four strains were excluded either because they were polyphyletic (SC-27) or had no NVT isolates present pre- or post-vaccine, and therefore could not be imputed (SC-04C, SC-12, and SC-17). The points are colored by serotype composition of strains: NVT only (blue) and mixed vaccine serotype (VT) and NVT (purple). The shaded quadrants indicate regions of accurate prediction of the prevalence change direction (increased post-vaccine vs. decreased) given the predicted fitness value. Three outlier strains are annotated for which the predicted direction of their prevalence change differed from that which was observed (i.e., they were predicted to increased based on their fitness when their prevalence from pre-to post-vaccine decreased, or vice-versa). **B.) Scatterplot of observed** versus **predicted prevalence of 27 strains at post-vaccine equilibrium based on quadratic programming**. These 27 strains contained at least one NVT strain pre-vaccine. Points are colored based on serotype composition as described in panel A. Perfect predictions would lie on dotted line of equality (1:1 line). The shaded grey region shows the confidence interval from the linear regression model used to test for deviation of the observed vs. predicted values compared to the 1:1 line. Two outliers are annotated for which the difference between their predicted and observed prevalence was >1.5 times the interquartile range of the distribution of predicted and observed prevalence differences. As a note, the predictions remained significant if SC-09 (the extreme strain at 10% prevalence in B) was removed (slope, 95% CI: 0.021, 1.05; intercept, 95% CI: −0.003, 0.03; p=0.19, chi-squared=3.5). **C-D.) Comparison of the predicted prevalence change from quadratic programming analysis using accessory genes and naïve *pro rata* model as shown in Figure 1B, but applied to just these 27 strains**. The dotted line of equality (1:1 line) and confidence interval (grey) are shown as in panel B. Goodness of fit statistics including sum of squared errors (SSE), root mean squared error (RMSE), and degrees of freedom adjusted R-squared (Adj. R^2^) are given for each model. The lower SSE and RMSE indicate a better model fit.

While the predicted fitness estimates how successful each strain will be immediately following vaccination, the long-term post-vaccine prevalence or change in prevalence of each strain is of more direct interest for evolution and public health. Thus, we posited that over time post-vaccination, gene frequencies would evolve to match as closely as possible to match those present pre-vaccination, and we used an optimization technique, quadratic programming, to calculate the NVT strain composition that produced accessory gene frequencies closest to those observed in the pre-vaccine population. Here we specifically focused on only the 27 strains that were observed pre-vaccine in the southwest US sample, allowing a projection with only data that was available at the time of vaccine introduction. This approach predicted the strain composition of the population following vaccination well, characterized by a 95% confidence interval of the observed vs. predicted post-vaccine strain frequencies that includes the line of equality (1:1 line), which denotes a perfect prediction, and by an intercept and slope that does not differ significantly from zero and one, respectively (p=0.24; intercept 95% CI: −0.005, 0.030; slope 95% CI: 0.257, 1.075, Figure 3B). Similar results were obtained when comparing predicted and observed change in prevalence (Figure 3C), where again the dotted line of equality fell within the 95% confidence interval of the regression of observed vs. predicted change in prevalence (p=0.75; intercept 95% CI: −0.02, 0.01; slope 95% CI: 0.23, 1.36). In comparison, a naïve *pro rata* estimate based solely on pre-vaccine prevalence performed poorly in predicting the prevalence change (Figure 3D, p=0.001; intercept 95% CI: −0.05, −0.008; slope 95% CI: −1.43, 0.35). In further support of these findings we examined a previously published carriage dataset of pneumococci colonizing children in Massachusetts. This dataset is imperfect in several respects. First, it was smaller (N=616), particularly the initial sample from the population, which had only 131 isolates and came in the first year of vaccine introduction rather than before it; we thus refer to it as “peri-vaccine.” Also making this data set less ideal, the last sample was obtained only six years after the first sample, giving less time for evolution to occur than in our southwest US data set. Changes in strain frequencies are shown in S2 Figure A-B. Despite the limitations of the data set, applying the same quadratic programming approach we could predict the post-vaccine equilibrium prevalence of the nine strains used in the analysis (p=0.65; intercept 95% CI: −0.05, 0.09; slope 95% CI: 0.25, 1.33) better than the *pro rata* model (S2 Figure C-E).

A further pneumococcal vaccine (PCV13) was introduced during the second half of our post-vaccine sampling of the southwest US dataset [17]. Despite this, the prevalence of PCV13 vaccine serotypes remained largely unchanged, suggesting little impact of this vaccine over the period of our study. To test the potential effect on our current analysis, we partitioned the post-vaccine sample into pre- and peri-PCV13 and the results are provided in Table 1, which demonstrate that our predictions were robust to sub-sampling. Finally, we tested the predictive value of different genomic elements, which are linked to accessory genes, finding that core genome loci (*n*_*loci*_= 17,101) and metabolic loci (*n*_*loci*_=5,853) were also capable of predicting the impact of vaccine, though not as accurately as the accessory genome based on goodness of fit statistics (Table 1). This finding must be considered in the context of recombination, selection, and the evolutionary timescale impacting the pneumococcal genome, which may impact the varying magnitude of NFDS signal across sets of loci. Despite moderate levels of bacterial recombination among pneumococci, there remains appreciable linkage disequilibrium between loci nearby as well as genome-wide [8], which makes it difficult to discern the relative selective importance of any particular locus. Exactly which genomic elements are responsible for the predictive ability we document here is unknown but is obviously of interest and should be a focus for future work.

**Table 1.** Comparison of pre-to post-vaccine prevalence change predictions using multiple models. Goodness of fit statistics including sum of squares due to error (SSE), root mean squared error (RMSE), and degrees of freedom adjusted R-squared (Adj. R^2^) are given for each model in relation to the 1:1 line. Fit statistics are provided for the naïve *pro rata* model and quadratic programming models using accessory genes, 5,853 biallelic polymorphic nucleotide sites found in 272 core-genome metabolic genes, and 17,101 biallelic polymorphic nucleotide sites found in 1,111 core genes. The results of the sensitivity analysis using a subsample of 119 isolates collected in 2010 prior to the initiation of PCV13 vaccine introduction is also presented for the accessory.

| Model | $n_{loci}$ | Adj. $R^2$ | SSE | RMSE |
| --- | --- | --- | --- | --- |
| *Pro-rata* (proportional change) | NA | 0.022 | 0.028 | 0.032 |
| Accessory genome (NFDS) | 2,371 | 0.223 | 0.015 | 0.024 |
| Accessory genome (NFDS) - Sensitivity analysis (2010 only) | 2,371 | 0.081 | 0.024 | 0.030 |
| Core genome (NFDS) | 17,101 | 0.173 | 0.016 | 0.024 |
| Metabolic loci (NFDS) | 5,853 | 0.154 | 0.017 | 0.025 |

In our main analysis, we can retrospectively calculate the predicted fitness of the two strains (shown as SC-10 and SC-24 in Figure 1 and S1 Figure) that emerged over the study period and compare them with contemporary samples collected elsewhere to determine their capacity for migration and emergence. Combining the southwest US dataset with the Massachusetts dataset [23,35], we identified 29 major strains and 2,511 accessory genes present between 5-95% among all 1,554 taxa. The predicted fitness values in our population after vaccine introduction range from −9.7 to 16.3 (median=2.5, SD=5.5) for strains present in the Massachusetts dataset. This included two strains, SC-10 (serotype 19A; ST320) and SC-24 (serotypes 15A, 23A/B; ST338), that were both present in the Massachusetts dataset peri-PCV7 (2001-2004) and also increased in prevalence thereafter. We found that these two strains had higher predicted fitness, 8.6 and 7.2 respectively for SC-10 and SC-24, than any of the other potential migrant strains that were not present in our southwest US sample before vaccination, indicating that their accessory gene frequencies were well adapted to offset the PCV7 perturbation in the southwest US population. Indeed, only two of the strains present before vaccination in the southwest US (SC-23 and SC-9) had a higher predicted fitness (Figure 3). This suggests we can use this approach to quantify which strains are most likely to successfully invade a population.

There are two primary ways in which NVT strains can fill the gap left in the population by vaccination, depending on their genomic relatedness to the removed PCV7 VT strains. First, NVT taxa that are closely related to VT strains in core and accessory genomes are opportune replacements and are therefore expected to by more successful than average following vaccination. There are two strains in our dataset that are exemplar of this, which both increased after vaccination (see SC-09 and SC-23 in Figure 1 and S1 Figure). We therefore expect that for any strain that contains both VT and NVT representatives, the NVT fraction will increase post vaccination, especially since these NVT taxa are sometimes similar to their VT counterparts in terms of serotype properties such as capsule thickness and charge, which are independently correlated with prevalence [36,37]. A good example of this is the serotype 15B/C component of strain SC-26 of the southwest US sample, which we now predict to be successful following the more recent introduction of a vaccine incorporating six additional serotypes (PCV13) and which has indeed been noted to be increasing in certain locations [38–40]. Second, where such close relatives are not available, the pre-vaccine frequencies of accessory genes can be restored by other NVT that are divergent in core genomes but similar in accessory genomes. This association likely often results from the movement of MGEs in the population (e.g., phages and transposons) or non-homologous recombination, which can make distantly related strains more similar in terms of genome content. As illustrated by the pairwise comparison of core/accessory genome divergence and absolute fitness difference of each strain (S3 Figure), there is an appreciable range of differences in fitness for strains that are equidistant in core and accessory genome divergence.

## Discussion

We show that by estimating the fitness of strains using an NFDS-based model and the frequencies of accessory genes, we are able to predict the direction of prevalence change following vaccination and more broadly the post-vaccine population composition. The ability of this type of balancing selection to determine the strain composition of a population is consistent with findings from environmental microbiology on multiple bacterial species [28]. Among pneumococci, changes in population dynamics after the introduction of vaccine have been explained by selection on many different aspects of the organism, including metabolic types, antibiotic resistance, carriage duration, recombination rates, and serotype competition, all of which are likely to be relevant contributors alongside, or components of, the accessory genome [30,31,41,42]. We provide a simple and effective approach for estimating the fitness of any strain in a population evolving under NFDS acting on accessory loci. All that is required is knowledge of the strain composition of the population and the accessory loci associated with each strain, as this approach does not depend on NFDS acting on particular known biological functions to predict the consequences of vaccination. It is quite conceivable that a minority of loci are involved, including even SNPs in the core genome, which also show a correlation (see Table 1 and [22]). We do not wish to imply that the sorts of selection discussed here act alone. Our previous work suggests the interplay between host immunity and polymorphic protein antigens may play a significant role [43], and other work suggests an important role for metabolic loci in the core genome [30]. Phage predation and defense as well as antibiotic resistance all likely contribute to the observed signal [21].

Certainly, as shown by outliers to predictions in Figure 3, we acknowledge that the model does not currently capture all population dynamics. Variation among loci in the strength of NFDS could account for some of these discrepancies, as indicated by retrospective model fitting. Other explanations include differences in the distribution of antibiotic resistance genes or possible vaccine cross-reactivity. For example, SC-18, containing serotype 6C, declined despite a positive predicted fitness; however, cross-reactivity between the PCV7 6B vaccine component and 6C may in part explain this observation [44]. Nevertheless, given the many potential pressures, mostly not directly observable, that we might expect to structure the pneumococcal population it is notable how effectively this approach can predict the impact of this perturbation. Overall, we find a significant relationship between predicted fitness and the adjusted prevalence change of a strain. By optimizing the prevalence of each strain conditional on the gene frequencies before vaccination we can estimate the equilibrium population after vaccination, using both the predicted fitness and numerical approximations of the post-vaccine equilibrium.

This work suggests numerous potential directions for future work, among them identifying the specific accessory loci or other genomic elements that are responsible for what we observe. Expanding the model to include immigration of other strains and disentangling the relative contribution of selection on various loci is likely to be a fruitful area for future research. One area worth exploring is the degree to which recombination acts to maintain gene frequencies on the timescale of population-level shifts in lineage composition. The emergence of new strains, characterized by novel combinations of accessory loci, is expected to be limited by the other strains present in the population in ways that are currently not well understood.

Predicting evolution is a central goal of population genomics especially when related to pathogens and human health. While evolutionary theory provides an understanding of bacterial population processes including the relative success of lineages, distribution of phenotypes, and ecological niche adaption, these analyses are often conducted retrospectively. Here, we demonstrate a method for predicting the impact of perturbing the pneumococcal population that may be useful to predict the outcomes of future interventions including vaccines. By incorporating information on invasive capacity, these predictions could be extended to inform changes in invasive disease rates. These dynamics may suggest novel vaccine strategies in which one could target those strains whose removal would result in a predicted re-equilibration that favors the least virulent or most drug-susceptible lineages [45]. The pervasive finding of accessory genomes in most bacterial species is usually explained by specialization of lineages to specific niches; however, it could also reflect widespread NFDS, and so future work should seek for evidence of similar signal in the core and accessory genome of other bacteria [46].

## Methods

### Study population and descriptive statistics (Figure 1 and S2 Figure)

The southwest US dataset used in this study is a subset of three studies of pneumococcal carriage conducted among Native American communities in the southwest US from 1998 to 2012, as previously described [47–49]. The pre-vaccine sample was collected from the well-defined control communities of the group-randomized trial of the PCV7 vaccine [48]. Pre-vaccine isolates included in our study were collected between March 1998 and April 2001. In late October 2000, PCV7 vaccination became routine, including catch-up for children aged <5 years. By March 2001, a total of 88% of 3–4-month old infants living in PCV7-randomized communities and 77% of those in control communities had received >1 dose of PCV7 [50]. However, only 7 of the 274 isolates in our pre-vaccine sample were collected between October 2000 and March 2001; therefore, we feel it is reasonable to treat it as a pre-vaccine sample from an unperturbed population. The 13-valent pneumococcal conjugate vaccine (PCV13) was later introduced in 2010. The pneumococcal sample was subdivided into 35 sequence clusters (SCs), referred to as strains in the main text, based on core genome diversity using hierarchical Bayesian Approximation of Population Structure (hierBAPS) [51]. Secondary strain clustering (e.g., A/B/C) was assigned using the second level clustering provided by hierBAPS analysis. A previously described carriage dataset of pneumococcal isolates from Massachusetts, US was also used to explore NFDS dynamics. For our analysis, we used the original population stratification of 16 strains identified by Croucher *et al*. [23,35]. For both datasets, we then classified strains by serotype composition as vaccine serotype (VT), non-vaccine serotype (NVT), or mixed (VT-NVT). The methods for whole-genome sequencing and genome assembly, and population genomic analysis have been described elsewhere [17].

For the present analysis, we focused on 937 pneumococcal carriage isolates from the southwest US collected during three study periods (epochs): pre-vaccine – population equilibrium (E1, 1998-2001); peri-PCV7 – population perturbation (E2, 2006-2008); post-PCV7 – population equilibration (E3, 2010-2012). The pre-vaccine period preceded the introduction of PCV7, while peri- and post-provided snapshots 5-6 and 10-12 years, respectively, after the introduction of PCV7. While the post-vaccine period includes, in part, the introduction of PCV13, we have previously shown that the majority of the sample was obtained when the impact of PCV13 was minimal [17]. This is supported by a sensitivity analysis to assess the effect of including all post-vaccine (E3) isolates by splitting sample into pre- and post-introduction and testing independently (2010 vs. 2011-2012).

For the additional dataset of carriage isolates from Massachusetts, we considered 133 isolates collected in 2001 as E1, even though the PCV7 was introduced in these communities in 2000, and 280 strains collected in 2007 as post-PCV7 (E3). Comparatively, the elapsed pre/post-vaccine time (E1-E3) differed considerably, being 6 years in the Massachusetts sample compared to 10-12 year in the southwest US sample. Based on previous analysis of the southwest US data, accessory gene frequencies were still experiencing perturbation 5-6 years after vaccine introduction (See S3 Figure in [17]). Therefore, it is likely that the Massachusetts sample had not yet reached a new post-vaccine equilibrium. We considered serotypes 4, 6A, 6B, 6E, 9V, 14, 18C, 19F, and 23F as PCV7 vaccine-type. For each strain, we computed the proportion of PCV7 VT and NVT. Three serogroup 6 serotypes were included because it has previously been shown that the serotype 6B component of PCV7 was cross-protective against 6A and that 6E produces a 6B capsular polysaccharide [52]. Further, cross-reactivity is consistent with the observed elimination of 6A and 6E in the study population after the introduction of PCV7 [17].

The observed changes in prevalence were estimated as 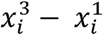, where 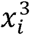 is the prevalence of strain *i* at E3 (post-vaccine) and 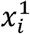 is the prevalence of strain *i* at E1 (pre-vaccine). As a null model for vaccine impact (*pro rata* model), we calculated the expected prevalence for each strain if its VT representation declined to zero in the whole population from pre-to post-vaccine, and its NVT representation increased proportionately to that in the whole population, and where the new NVT prevalence values *g*_*i*_ are renormalized to sum to one. We defined the prevalence change as 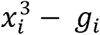. To determine significant deviations of the observed post-vaccine strain prevalence from the *pro rata* model, we sampled 10,000 bootstrap replicates with replacement from E1, and calculated the *pro rata* prevalence changes for each replicate. We then plotted the 2.5%, 50%, and 97.5% quantiles of these resampled predictions in Figure 1B. We defined 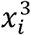 as significantly different from the null expectation if the strain’s prevalence change was outside the central 95% of the bootstrap distribution of the predicted value.

### Pneumococcal pangenome analysis

As previously described, pangenome analysis of 937 taxa was carried out using Roary v3.12.0 [17]. The resulting presence/absence matrix was used to generate a binary accessory genome alignment of 2,371 clusters of orthologous groups (COGs). This binary alignment was used to infer a maximum likelihood (ML) phylogeny using RAxML v8.2 with BINGAMMA substitution model and 100 bootstrap replicates [53]. The same approach was used to infer a ML phylogeny of SNPs found in the core genome using the GTRGAMMA substitution model. Serotype, collection period (epoch), and strain (SC) assignment were visualized in relation to the accessory genome phylogeny. We then imported the phylogeny into R using APE v4.1 and computed the mean pairwise patristic distance among all strains using the *meandist* function in the R package Vegan v2.4-67 [54]. Hierarchical clustering of scaled between-strain patristic distances was visualized using *heatmap*.*2* in ggplots v3.0.1. Last, core and accessory genome divergence was compared to the absolute fitness difference among strains. For the additional carriage dataset from Massachusetts, the presence/absence matrix was obtained from the online repository available at https://www.nature.com/articles/sdata201558.

### Predicted Fitness

In the southwest US dataset, we identified 35 strains among 937 isolates. This included a polyphyletic grouping of strains present at low frequencies in the overall population (SC-27). Pre-vaccine, two strains (SC-10 and SC-24) were not sampled, having only been observed after the introduction of vaccine. Further, two strains (SC-22 and SC-23) had no NVT component pre-vaccine but did post-vaccine. For these four strains, we imputed pre-vaccine accessory gene frequencies by subsampling representative taxa from the first time point when they were observed (peri-vaccine period (E2) in both instances). This allowed us to calculate the fitness of these strains. Three additional strains (SC-04C, SC-12, and SC-17) were excluded because they had no NVT isolates present pre-vaccine or were not observed post-vaccine (i.e., they were comprised solely of VT isolates); therefore, their fitness could not be imputed nor their prevalence change. Finally, there were a few instances of strains that contained both VT and NVT serotypes. Where this was the case, for the purposes of considering the NVT portion of such strains, we removed the VTs and considered the remainder in isolation as an NVT strain. This was repeated for 14 of 16 strains in the carriage dataset from Massachusetts. This required imputing five strains that were not sampled pre-vaccine.

For the two previously unobserved strains (SC-10 and SC-24) in the primary dataset, we assessed the degree to which their accessory genome composition may have contributed to emergence after the introduction of PCV7 by comparing their fitness to strains found in the Massachusetts dataset [23,35]. To do this, we repeated the pangenome analysis using a merged dataset of 1,554 carriage isolates (including all genomes from [24]). Population structure (determination of strains) of the combined sample was assessed with hierBAPS and accessory gene filtering was conducted as previously detailed. Frequencies of accessory genes were determined for each strain in the Massachusetts dataset, and the predicted fitness values were calculated by comparing those frequencies to (*e* − *f*) in the primary southwest USA dataset. The distribution of fitness values in the Massachusetts dataset were assessed and compared with the two emergent strains to determine their ranking. Last, to predict the impact of PCV13 on the pneumococcal population, we repeated the quadratic programming analysis on the post-vaccine population. To do this, we recalculated the change in strain prevalence resulting from the removal of six additional PCV13 VT serotypes (1, 3, 5, 6A, 7F, 19A) and determined the predicted fitness for each extant NVT strain to identify those with positive values, i.e. those that will likely be more successful in the PCV13 era.

### Post-vaccine equilibrium frequencies via quadratic programming

Using 2,371 accessory genes present in 5-95% of taxa of the southwest US dataset, we determined pre-vaccine accessory gene frequencies for each strain, considering NVT taxa only. For this, we focused on 27 major strains which 1) had NVT taxa present pre-vaccine and 2) were not polyphyletic. This excluded eight strains (SC-04C, SC-10, SC-12, SC-17, SC-22, SC-23, SC-24, and SC-27) and replicated what would have been possible with the available pre-vaccine data. S4 Figure shows the distribution of the 2,371 accessory genes among isolates belonging to the 27 strains. This figure was also used to test the assumption that the impact of recombination on the accessory genome is negligible over our study period, where we compared the pre-vaccine and post-vaccine accessory gene frequencies for each NVT strain. For the 27 strains, we computed the predicted prevalence of each strain such that post-vaccine accessory gene frequencies approached as closely as possible to pre-vaccine frequencies by using a quadratic programming approach. Quadratic programming involves optimizing a quadratic function based on several linearly constrained variables [55], and was done using the package quadprog v1.5-5 implemented in Rstudio v1.0.143 with R v3.3.19 [56]. Details of this implementation can be found in the R code provided. This was then repeated using: 1) 17,101 biallelic polymorphic sites found in 1,111 genes in the core genome and present among 5-95% of taxa and 2) 5,853 biallelic polymorphic sites found in 272 metabolic genes present in the core genome and present among 5-95% of taxa. We then conducted a sensitivity analysis using genes present in 1-99% and 2.5-97.5% of taxa and found the results did not differ significantly from those obtained using genes present among 5-95% of taxa. Detailed methods for the ascertainment of genomic loci are in Azarian *et al* [17].

Using 1,056 accessory genes present in 5-95% of taxa of the Massachusetts dataset, we determined pre-vaccine accessory gene frequencies for each strain, considering NVT taxa only. For this, we focused on 9 major strains, which had NVT taxa present pre-vaccine and were not polyphyletic (SC-1, SC-2, SC-4, SC-8, SC-9, SC-10, SC-11, SC-12, SC-16). This excluded seven strains and replicated what would have been possible with the available pre-vaccine data. For the 9 strains, we computed the predicted prevalence of each strain such that post-vaccine accessory gene frequencies approached as closely as possible to pre-vaccine frequencies using a quadratic programming as described above.

For each model, we evaluated accuracy by determining if the slope and intercept of the predicted and observed strain frequencies were close to one and zero, respectively. Goodness of fit statistics including sum of squares due to error (SSE), root mean squared error (RMSE), and degrees of freedom adjusted R-squared (Adj. R^2^) were used to evaluate each model. In addition to assessing how well we could predict post-PCV7 prevalence, we also tested if we accurately inferred whether a strain would increase or decrease after the introduction of vaccine. To do this, we calculated the observed prevalence trajectory from pre-to post-vaccine and compared that to the predicted trajectory, identifying those with significantly positive or negative risk differences using Fisher’s exact test.

## Data and materials availability

Whole-genome sequencing data are available from NCBI under BioProject PRJEB8327: https://www.ncbi.nlm.nih.gov/bioproject/PRJEB8327. Accession numbers and accompanying metadata have previously been published. All R code and associated input data frames as well as phylogenies are available on GitHub here: https://github.com/c2-d2/Projects/tree/master/NFDS.

## Acknowledgements

The ideas in this paper and the collaborative relationships underlying them are owed in significant part to the Permafrost Conference on Microbial Genomics, of Blessed Memory. We would also like to thank Xueting Qiu for her assistance with the manuscript preparations and revisions.

