## Supplemental figures for "Predicting evolution using frequency-dependent selection in bacterial populations"

**1** Burnett School of Biomedical Sciences, University of Central Florida, Orlando, FL; **2** Center for Communicable Disease Dynamics, Department of Epidemiology, T.H. Chan School of Public Health, Harvard University, Boston MA; **3** Center for American Indian Health, Johns Hopkins Bloomberg School of Public Health, Baltimore, Maryland; **4** Helsinki Institute for Information Technology, Department of Mathematics and Statistics, University of Helsinki, 00014 Helsinki, Finland. **5** Department of Biostatistics, University of Oslo, 0317 Oslo, Norway;

**6** Infection Genomics, The Wellcome Trust Sanger Institute, Wellcome Trust Genome Campus, Hinxton, Cambridge CB10 1SA, UK; **7** Big Data Institute, Nuffield Department of Medicine, University of Oxford, Oxford OX3 7LF, UK; **8** MRC Centre for Global Infectious Disease Analysis, Department of Infectious Disease Epidemiology, Imperial College London, London W2 1PG, UK; **9** World Health Organization, Geneva Switzerland; **10** Department of Immunology and Infectious Diseases, T.H. Chan School of Public Health, Harvard University, Boston MA.

Pamela P Martinez

Brian J Arnold

Lindsay R Grant

Jukka Corander

Christophe Fraser

Nicholas J Croucher

Laura Hammitt

Raymond Reid

Mathuram Santosham

Robert R Weatherholtz

Stephen D Bentley

Katherine L O’Brien

Marc Lipsitch

William P Hanage

**Corresponding Author:*

Taj Azarian, PhD MPH

Burnett School of Biomedical Science, College of Medicine

University of Central Florida

**Supplement figures:**

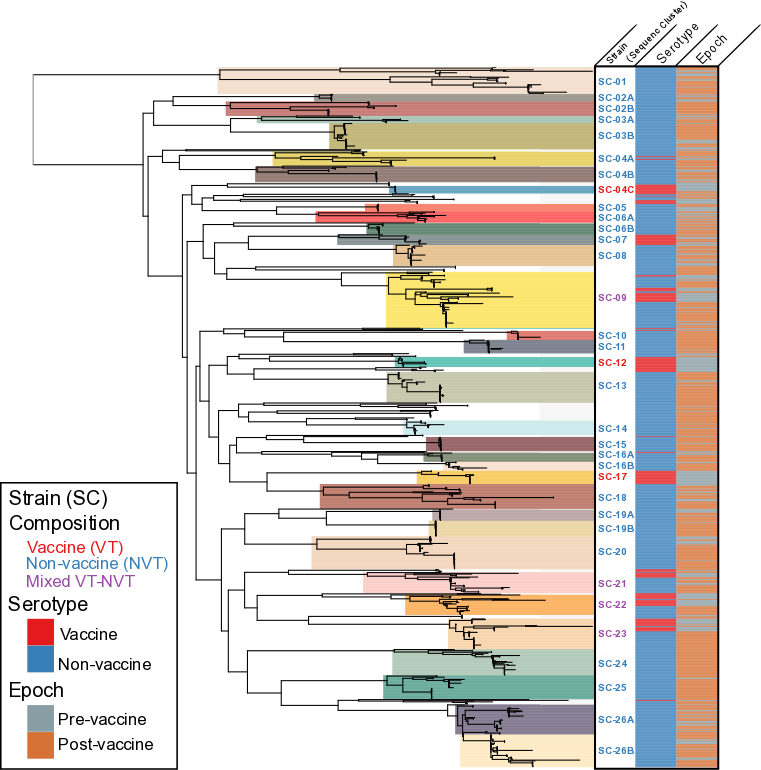

**S1 Figure.** Maximum likelihood phylogeny inferred from an alignment of single-nucleotide polymorphisms present in the core genome of 937 isolates obtained from a subset of three studies of pneumococcal carriage conducted among Native American communities in the southwest US from 1998 to 2012. The collection epoch, pre-vaccine or peri/post-vaccine, is indicated by the grey or orange. Thirty-five strains (i.e., sequence clusters, SCs), which were identified by subdividing the pneumococcal population on core genome diversity, are shaded by color on the phylogeny and labeled to the right. Sub-clusters (e.g., A/B/C) were identified using the second-order clustering of BAPS analysis. The phylogeny is ordered by branch decreasing branch length, and strains are numbered such that closer numbers are more genomically similar. The text color of the strain label indicates the serotype composition. The heatmap indicates PCV7 vaccine (VT) and non-vaccine (NVT) serotypes and collection period, illustrating the removal of vaccine serotypes by the introduction of PCV7. As described in the text, two NVT strains were not present pre-vaccine but subsequently increased thereafter.

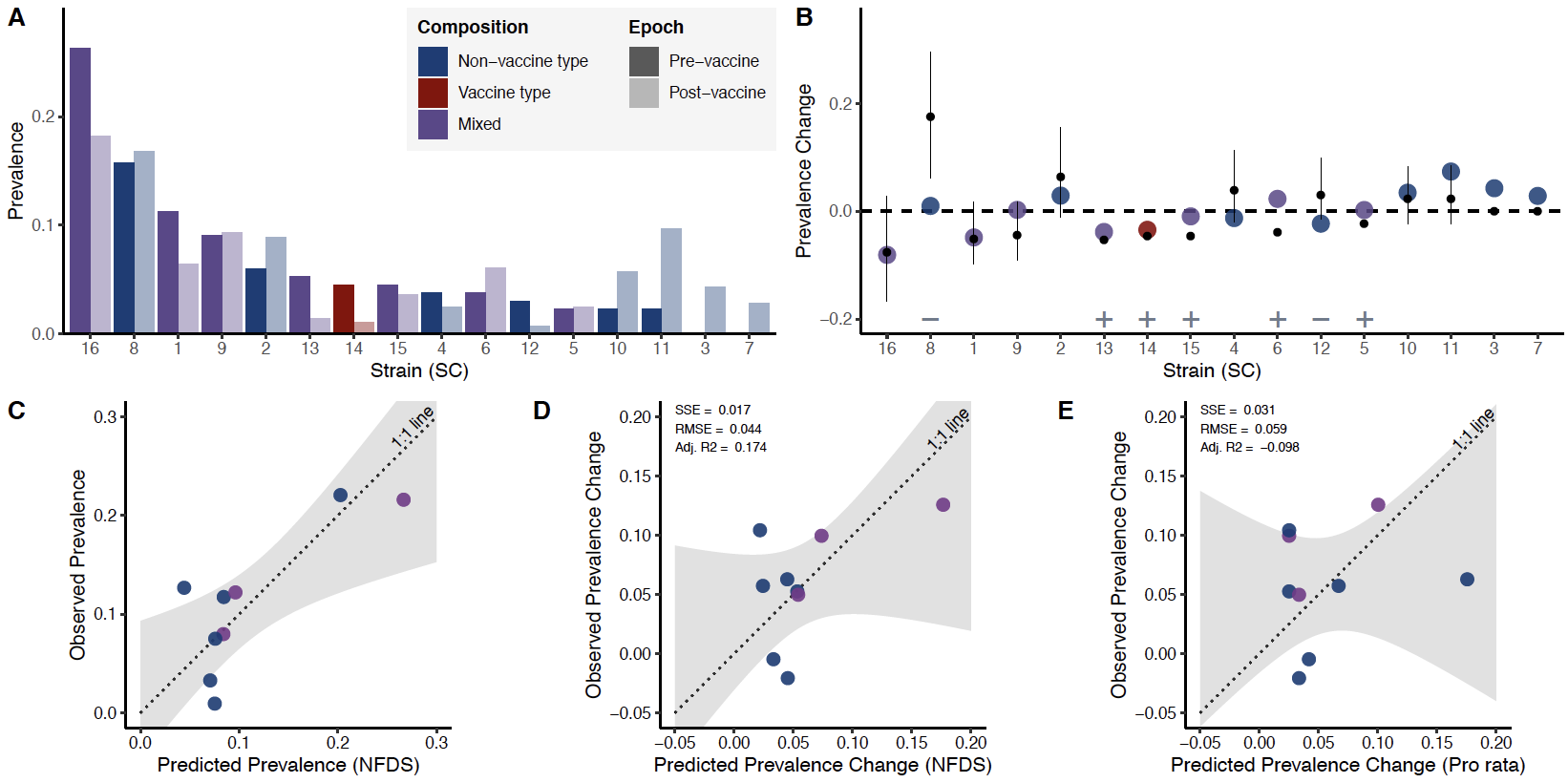

**S2 Figure**. Analysis of 616 pneumococcal carriage isolates collected from Massachusetts children. A.) Peri-vaccine (E1, 2001) to post-vaccine (E3, 2007) change in prevalence of 16 strains (sequence clusters, SCs). Strains are ordered from highest to lowest pre-vaccine prevalence. B.) Change in prevalence from peri-vaccine (E1) to post-vaccine (E3) ordered by strain as in (A). Observed changes in prevalence are represented by points colored by serotype composition of the strain: non-vaccine serotype (NVT) only, PCV7 vaccine-serotype (VT) only, and mixed VT and NVT (VT-NVT). The point and whiskers show the prevalence change expected if all VT strains were removed and NVT increased *pro rata* to their pre-vaccine prevalence. The dot is the median, and the whiskers give the 2.5% and 97.5% quantiles of predicted changes under the null model using 10,000 bootstraps from pre-vaccine and post-vaccine samples. Significant differences are denoted with plus and minus signs specifying strains that were significantly more (n=5) or less (n=2) common, respectively, than expected under the null model. C.) Scatterplot of observed versus predicted prevalence of nine strains at post-vaccine equilibrium based on quadratic programming. These nine strains contained at least one NVT strain pre-vaccine. Points are colored based on serotype composition as described in panel A. Perfect predictions would lie on the dotted line of equality (1:1 line). The shaded grey region shows the confidence interval from the linear regression model used to test for deviation of the observed vs. predicted values compared to the 1:1 line. D-E.) Comparison of the predicted prevalence change from quadratic programming analysis using accessory genes (D, p=0.38; intercept 95% CI: -0.08, 0.03; slope 95% CI: -0.24, 1.32) and naïve *pro rata* model (E, p=0.07; intercept 95% CI: -0.11, 0.02; slope 95% CI: -0.65, 1.03) as shown in panel A. Goodness of fit statistics including sum of squares do to error (SSE), root mean squared error (RMSE), and degrees of freedom adjusted R-squared (Adj. R2) are given for each model. The lower SSE indicates a better model fit. Panel A presents the frequencies of 16 strains based on the entire population, while the analysis for C, D, and E necessitated the use of frequencies based on nine strains.

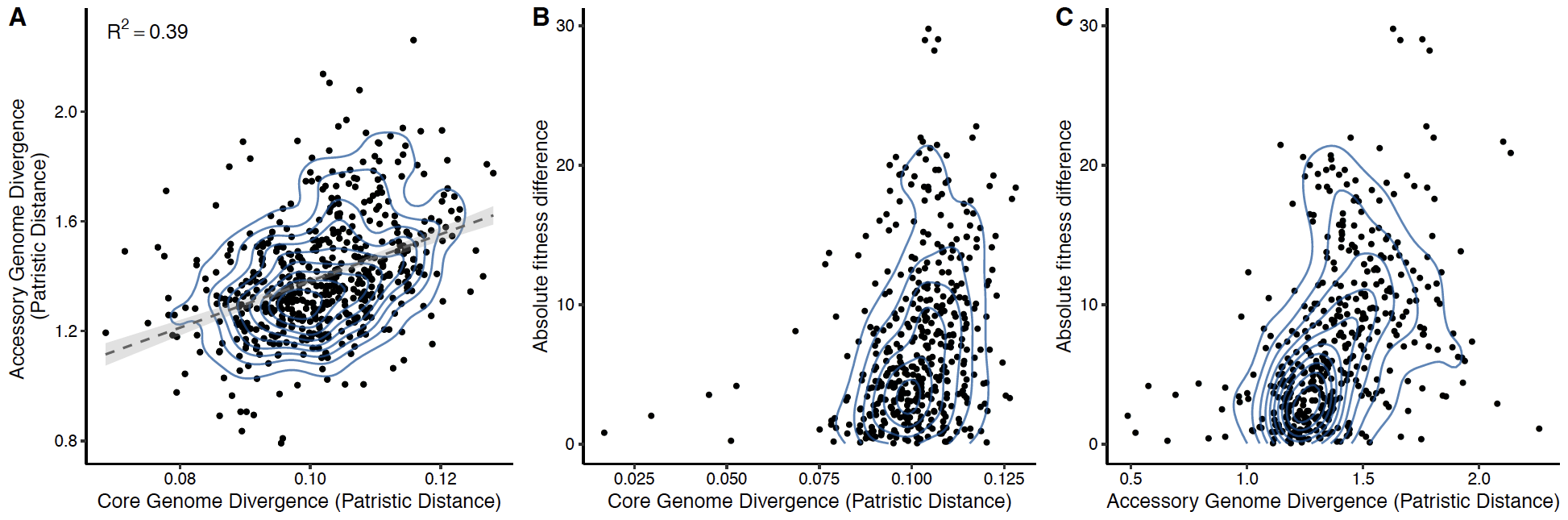

**S3 Figure.** **A.) Comparison of core and accessory genome divergence, quantified as the mean patristic distance between strains (sequence clusters, SCs) on respective core and accessory genome maximum likelihood phylogenies.** Each point represents a pairwise comparison of the mean between strain patristic distances. Pairwise comparisons between strain sub-clusters (e.g., SC-25A and SC-25B, see S1 Figure phylogeny) and between the unencapsulated non-typeable SC-01 and other strains have been excluded. This removes the most and least divergent pairwise comparisons, which otherwise artificially inflated the linear relationship between core and accessory genome distances. The dotted line represents the fit of a linear regression model to the data with 95% confidence interval shaded in grey. Contour lines show two-dimensional kernel density estimation based on the distribution of points. **B.) Core genome divergence (as defined above based on the core genomic phylogeny patristic distances) and the absolute fitness difference among 31 strains presented in Figure 3A. C.) Accessory of core genome divergence (as defined above based on the core genomic phylogeny patristic distances) and the value of the fitness difference among 31 strains presented in Figure 3A**. Of note, there is a considerable range in predicted fitness difference among strains that have similar accessory and core genome divergence (e.g., core genome divergence of 0.1 or accessory genome divergence of 1.0-1.5).

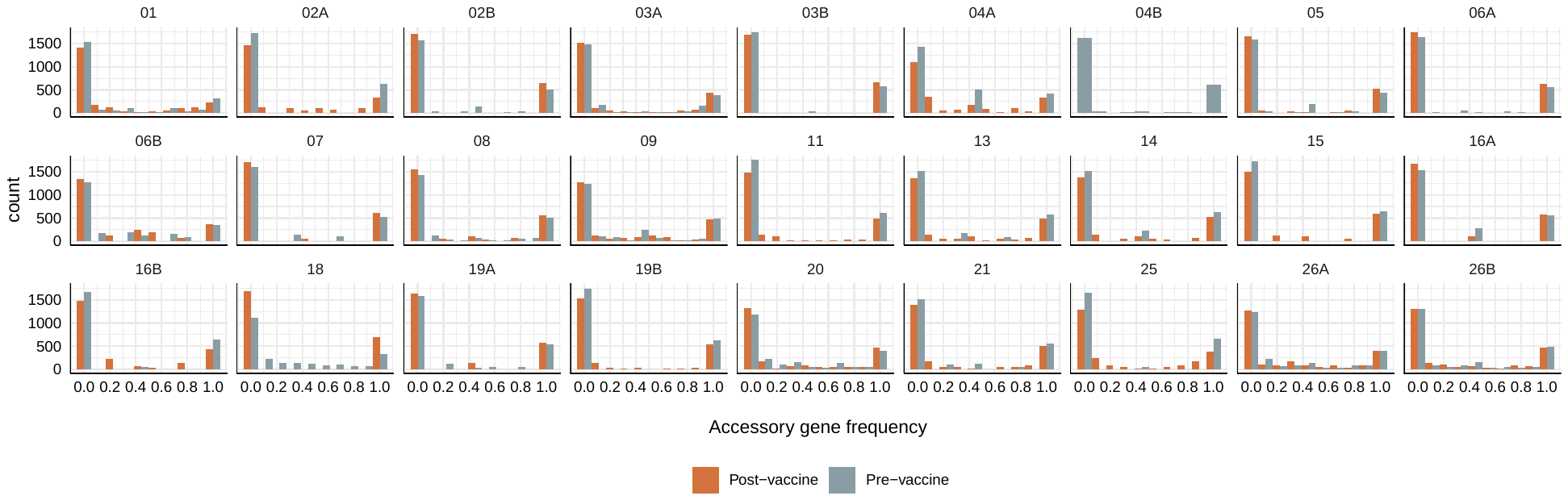

**S4 Figure.** Distribution of clusters of orthologous groups among 27 non-vaccine type (NVT) strains that were observed pre-vaccine, colored by the epoch. Histograms represent the presence and absence of accessory genes that were present in between 5% and 95% of the entire sample. Note, each strain will have a variable number of accessory genes depending on their accessory genome diversity. Within each strain, the distribution of accessory genes remained relatively stable from pre- to post-vaccine.
